# CATaDa reveals global remodelling of chromatin accessibility during stem cell differentiation in vivo

**DOI:** 10.1101/189456

**Authors:** Gabriel N. Aughey, Alicia Estacio Gomez, Jamie Thomson, Hang Yin, Tony D. Southall

**Affiliations:** Department of Life Sciences, Imperial College London, Sir Ernst Chain Building, London, UK

## Abstract

Regulation of eukaryotic gene expression is coordinated by dynamic changes to chromatin states throughout development. Measurements of accessible chromatin are used extensively to identify genomic regulatory elements. Whilst the chromatin landscapes of pluripotent stem cells are well characterised, chromatin accessibility changes in the development of somatic stem cell lineages are not well defined. Here we show that tissue specific chromatin accessibility data can be produced via ectopic expression of *E. coli* Dam methylase *in vivo*, without the requirement for cell-sorting. We have profiled chromatin accessibility in individual cell types of the *Drosophila* neural and midgut stem cell lineages. Functional cell-type specific enhancers were identified, as well as novel motifs enriched at different stages of development. Finally, we show global changes in the accessibility of chromatin between stem-cells and their differentiated progeny. Our results demonstrate the dynamic nature of chromatin accessibility in somatic tissues during stem cell differentiation and provide a novel approach to understanding the gene regulatory mechanisms underlying development.

## Introduction

During the development of a multicellular organism, gene expression is tightly regulated in response to spatially and temporally restricted signals. Changes to gene expression are accompanied by concomitant changes to chromatin structure and composition. Therefore chromatin states vary widely across developmental stages and cell types. Functional regions of a genome, including promoters and enhancers, can be identified by their relative lack of nucleosomes. These regions of “open chromatin” can be assayed by their accessibility to extrinsic factors. Consequently, chromatin accessibility profiling techniques are commonly used to investigate chromatin states (reviewed in [1]). Chromatin is highly accessible in pluripotent cell types such as embryonic stem (ES) cells, but is compacted following differentiation [2]. It has been suggested that this open chromatin represents a permissive state to which multiple programmes of gene regulation may be rapidly applied upon differentiation [3].

The nature of chromatin accessibility across different developmental stages *in vivo* is less well understood. Imaging studies have been used to demonstrate gross changes to chromatin structure, for example changes to the distribution of heterochromatin have been observed in post-mitotic cells [4, 5]. Molecular studies investigating chromatin states *in vivo* during development have tended to utilise heterogeneous tissues due to the fact that profiling the epigenome of individual cell types frequently requires physical isolation of cells or nuclei, which can be laborious and prone to human error [6]. Therefore, there is a lack of information regarding cell-type specific changes to chromatin states in *in vivo* models. Whilst recently developed methods such as ATAC-seq have become popular and address many of the limitations inherent to earlier techniques such as DNAse-seq (i.e. requires fewer cells and increased assay speed), these techniques still require the physical separation of cells and isolation of genomic DNA before chromatin accessibility is assayed [7].

It has been suggested that ectopic expression of untethered DNA adenine methyltransferase (Dam) results in specific methylation of open chromatin regions whilst nucleosome bound DNA is protected [8-11]. However, the efficacy of using Dam methylation for chromatin accessibility profiling on a genomic scale is not clear. Furthermore, expression of Dam in a cell-type specific manner, at levels low enough to avoid toxicity and oversaturated signal, has not been possible until now.

Transgenic expression of fusions of Dam to DNA binding proteins is a well-established method used to assess transcription factor occupancy (DNA adenine methyltransferase identification - DamID) [12]. Recently it was demonstrated that DamID could be adapted to profile DNA-protein interactions in a cell type specific manner by utilising ribosome re-initiation to attenuate transgene expression [13-15]. This technique is referred to as Targeted DamID (TaDa). Here we take advantage of TaDa to express untethered Dam in specific cell-types to produce chromatin accessibility profiles *in vivo*, without the requirement for cell separation. We show that **C**hromatin **A**ccessibility profiling using **Ta**rgeted **Da**mID (CATaDa) yields comparable results to both FAIRE and ATAC-seq methods, indicating that it is a reliable and reproducible method for investigating chromatin states. By assaying multiple cell types within a tissue, we show that chromatin accessibility is dynamic throughout the development of *Drosophila* central nervous system (CNS) and midgut lineages. These data have also enabled us to identify enriched motifs from regulatory elements that dynamically change their accessibility during differentiation, as well as to identify functional cell-type specific enhancers. Finally, we show that compared to their differentiated progeny, somatic stem cell Dam-methylation signals are more widely distributed across the genome, indicating a greater level of global chromatin accessibility.

## Results

### CATaDa produces chromatin accessibility profiles comparable to that of ATAC and FAIRE-seq in Drosophila eye discs

We reasoned that low-level expression of transgenic *E. coli* Dam, using tissue specific GAL4 drivers in *Drosophila*, would specifically methylate regions of accessible chromatin exclusively in a cell-type of interest. Detection of these methylated sequences could yield chromatin accessibility profiles for defined cell populations *in vivo* (Figure 1). To determine if CATaDa produces an accurate reflection of chromatin accessibility, we compared data acquired using this approach with commonly used alternative techniques. A recent study generated ATAC and FAIRE-seq data from *Drosophila* imaginal eye discs [16]. Using CATaDa, we expressed *E. coli Dam* in the eye disc of *Drosophila* third instar larvae so that we could compare *Dam* methylation profiles to these previously collected data.

**Figure 1.**
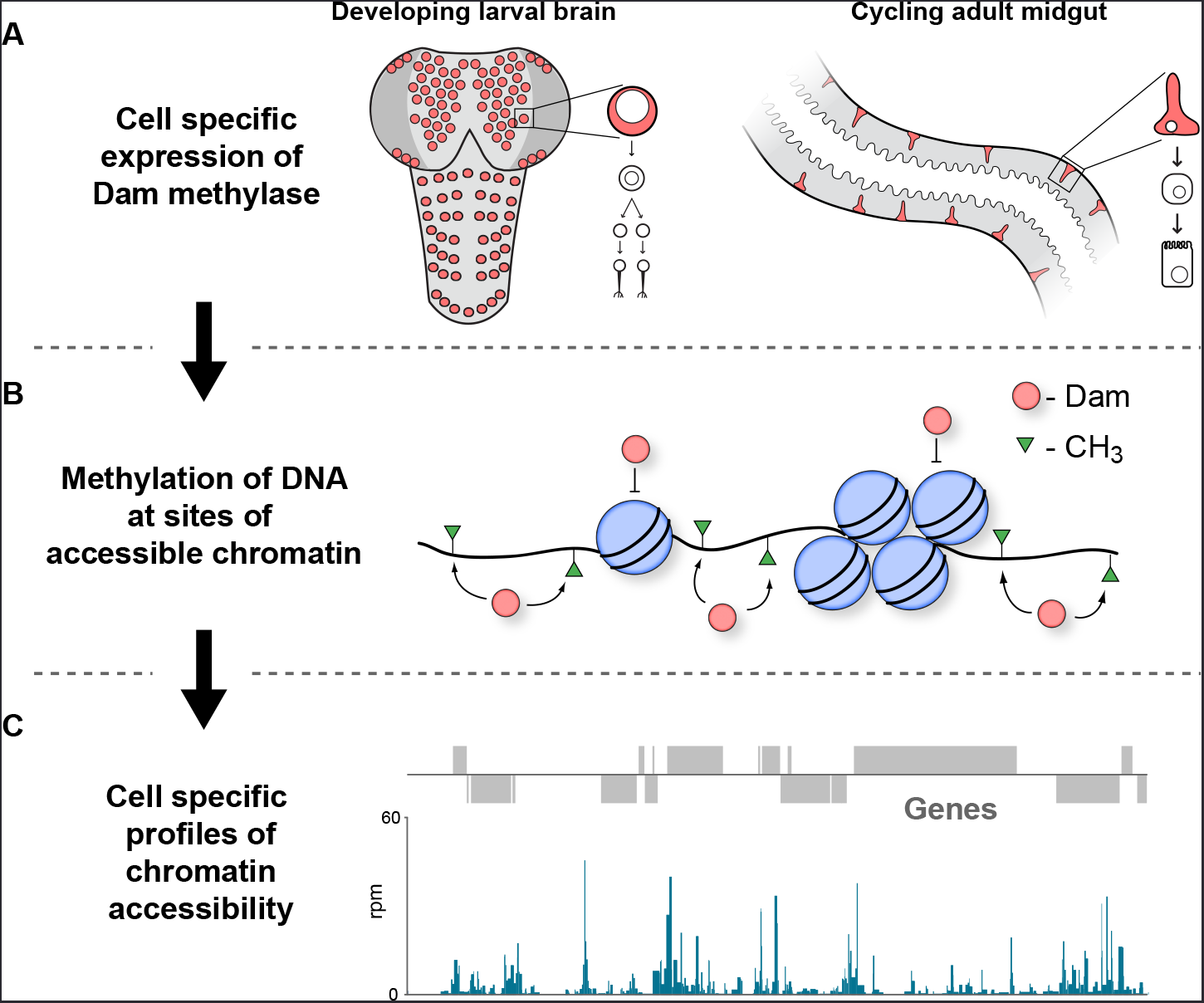
Schematic illustrating CATaDa technique. **A)** *E. coli* Dam is expressed specifically in cell-types of interest using TaDa technique. **B)** GATC motifs in regions of accessible chromatin are methylated by Dam, whilst areas of condensed chromatin prevent access to Dam thereby precluding methylation. **C)** Methylated DNA is detected to produce chromatin accessibility profiles for individual cell-types of interest from a mixed population of cells.

Chromatin accessibility profiles produced with CATaDa are highly reproducible between replicates (r^2^ = 0.947) (Supplementary figure 1). CATaDa profiles showed good agreement with data produced with ATAC-seq and FAIRE-seq. Visual inspection of the data showed that many regions of accessible chromatin identified by ATAC and FAIRE are also represented by CATaDa, whilst condensed regions are reliably inaccessible (Figures 2A,B). The overlap of Dam identified peaks with ATAC and FAIRE peaks is 48.5% and 49.4% respectively (In comparison, 64.8% of ATAC peaks are also identified in FAIRE data). A Monte Carlo simulation determined that this is a highly significant overlap (p< 1x10^-5^). We found that CATaDa profiles exhibited features consistent with chromatin accessibility. For example, open chromatin is enriched at transcriptional start sites (TSS) (Figure 2C). This is despite the relative depletion of GATC sites around TSS (Supplementary figure 2). It was previously shown that ATAC-seq and FAIRE-seq data demonstrated high chromatin accessibility at experimentally validated eye-antennal enhancers. CATaDa profiles similarly showed increased open chromatin at these regions (Figure 2B,D). We found that for 57.9% of FlyLight eye enhancers, a corresponding peak was called in CATaDa profiles (333 of 575 enhancers). CATaDa was comparable to FAIRE-seq and ATAC-seq which identified 48% and 68.7% respectively, of validated FlyLight enhancers as peaks (Figure 2E).

**Figure 2.**
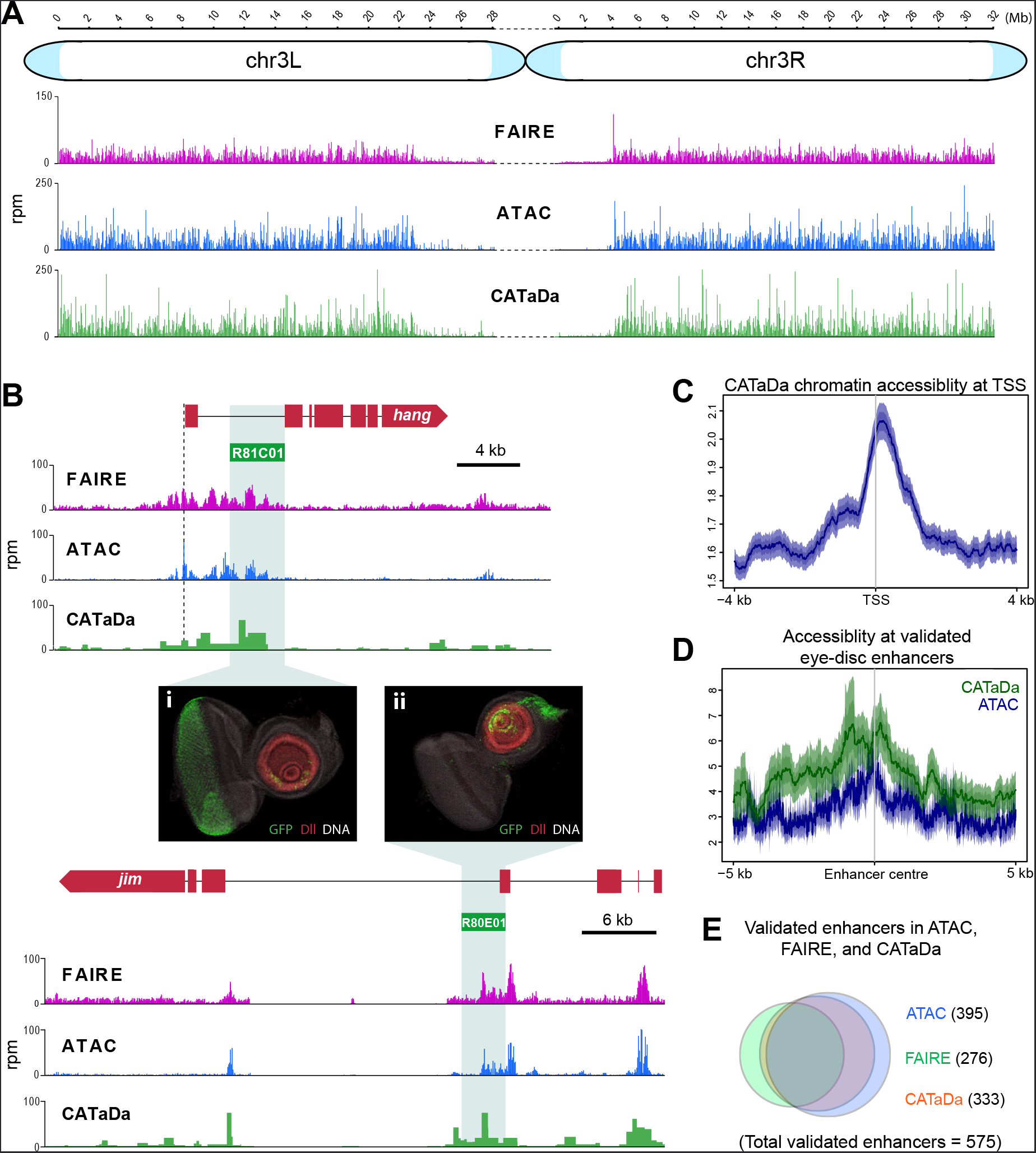
Validation of Dam chromatin accessibility profiling compared to ATAC and FAIRE-seq. **A)** Chromatin accessibility across chromosome three as determined by ATAC-seq, FAIRE-seq, and CATaDa. Note the reduced amount open chromatin proximal to the centromere regions in all three datasets. y-axes = reads per million (rpm). **B)** Example loci showing data obtained by FAIRE, ATAC, and Dam accessibility profiling. Peaks are broadly reproducible across techniques. Flylight enhancers with validated expression in eye imaginal discs coincide with peaks in all three datasets. Corresponding expression pattern is shown in i) and ii) (eye disc images obtained from the FlyLight database (http://flweb.janelia.org/cgi-bin/flew.cgi)). **C)** Aggregation plot of Dam signal at TSS with 2 kb regions up and downstream. Aggregated signal at TSS shows expected enrichment of Dam, although signal is less strong than that of ATAC shown in (D). **D)** Aggregation plot showing average signal of ATAC (blue) and Dam (green) at 575 FlyLight enhancers with validated eye imaginal disc expression. Both techniques show increased open chromatin at these regions. **E)** Venn diagram of FlyLight enhancers identified in Dam accessibility profiling, ATAC, or FAIRE-seq. The majority of enhancers identified by either ATAC or FAIRE are also found in the Dam data. Dam enhancers overlap most with ATAC (305 shared between ATAC and Dam of 575 total FlyLight enhancers).

### CATaDa profiling shows dynamic changes in chromatin accessibility during differentiation of the nervous system

In *Drosophila*, neurons are derived from asymmetrically dividing neural stem cells (NSCs). NSC divisions produce one self-renewing daughter NSC and a ganglion mother cell (GMC), which divides once more to produce neurons or glia [17]. To investigate how local and global chromatin accessibility changes during the process of nervous system differentiation, we expressed Dam in specific cells with GAL4 drivers that cover four different developmental stages within the lineage. These include NSCs (*worniu-GAL4*), GMCs and newly born neurons (*R71C09-GAL4* (see Supplementary figure 4)[18]), differentiated larval neurons (nSyb-GAL4), and also mature adult neurons (*nSyb-GAL4*) (Figure 3A).

By examining candidate genes differentially expressed during neural development, we observed that chromatin accessibility relates to gene expression in an expected manner. For example, intronic open chromatin peaks can be seen at the *bruchpilot* (*brp*) locus, in both third instar (L3) and adult neurons, whilst these peaks are reduced or absent in the progenitor cell types (Figure 3B). This corresponds with the expression of *brp*, which is specifically transcribed in neurons and has an important role in synapse function [19]. In contrast, the adjacent gene to *brp, Wnt2*, displays peaks which are most apparent in the NSC and intermediate cell types. Wnt signalling is known to be important for the control of stem cell populations, therefore these results are also expected [20].

Similar patterns are observed at a number of other loci. At the *asense* (*ase*) locus, (a NSC-specific transcription factor), chromatin is highly accessible at the promoter and upstream intergenic region in NSCs (Supplementary figure 3). This signal is considerably reduced in fully differentiated neurons in which ase is not expressed. Interestingly, open chromatin is still detectable in these regions in the GMCs/ newly born neurons. This pattern is also observed with other NSC expressed factors such as *deadpan* (*dpn*), *CyclinE* (*CycE*) and *prospero* (*pros*) (Supplementary figure 3). Interestingly, GMC/ newly born neuron profiles frequently show intermediate signal at these loci, indicating that functional elements required for regulation of NSC gene expression are not immediately rendered inaccessible following differentiation (Figure 3B and supplementary figure 3).

**Figure 3.**
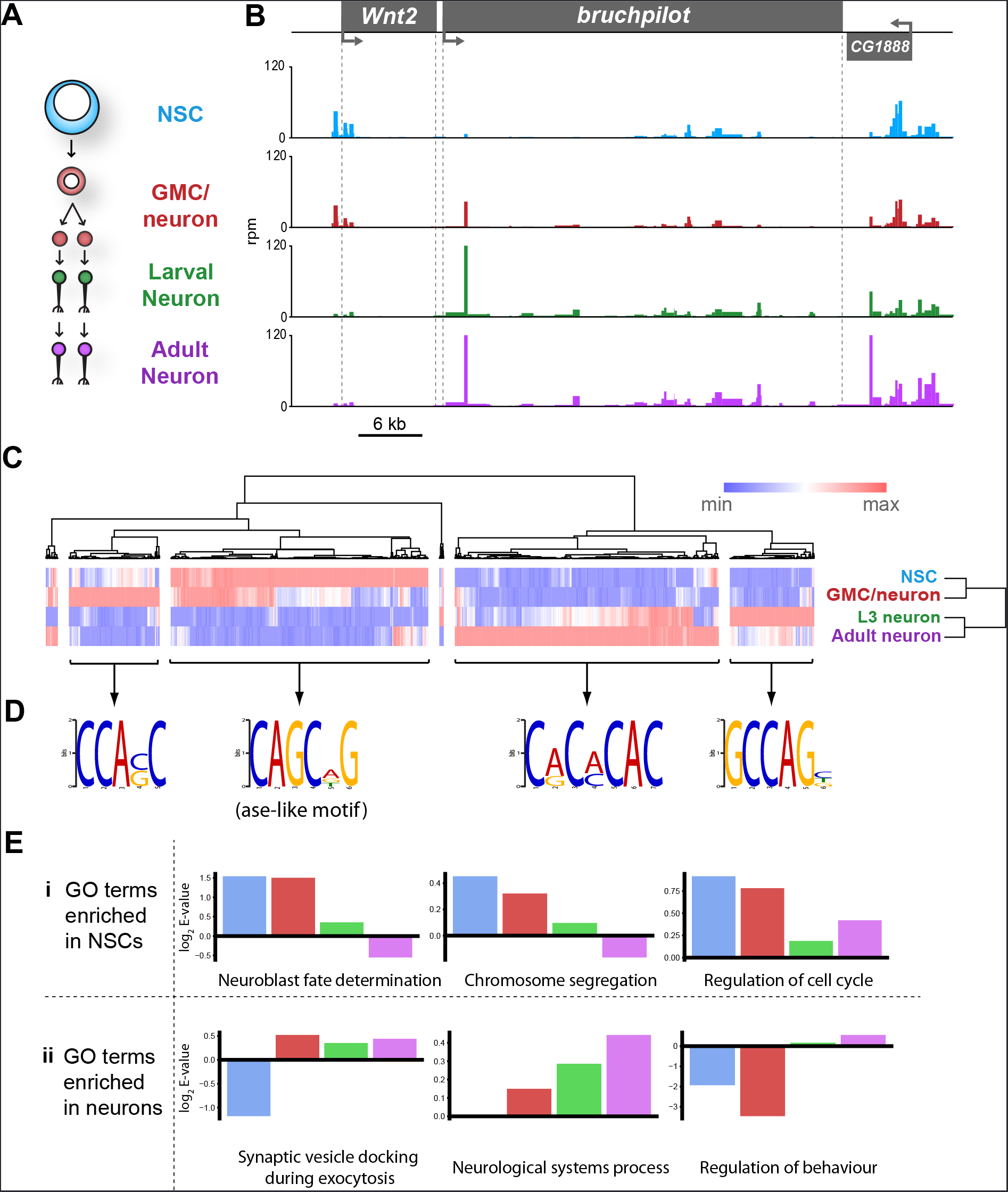
Chromatin accessibility of cell types in the CNS. **A)** Schematic of CNS lineage progression indicating cell types examined in this study. **B)** Example profiles resulting from Dam expression in the CNS. Genomic region encompassing Wnt2 and bruchpilot genes is shown. Multiple open chromatin regions are dynamic across development. Y-axes = reads per million (rpm). **C)** Clustering of differentially accessible regions in CNS lineages indicates two major groupings in which chromatin is most accessible in either stem cells or mature neurons. **D)** Motif analysis using these sequences results in identification of expected motifs (e.g. ase E-box motif in stem cell accessible loci), as well as novel motifs. Most highly enriched motifs for each cluster shown. All motifs E-values < 1x10^-5^. **E)** log2 enrichment scores for selected GO terms in individual cell types. Clear trends can be seen as development progresses. (NSC, GMC, L3 neuron, adult neuron - from left to right). **i)** GO terms are either enriched in stem cells becoming less significant as the lineage progresses or **ii)** vice versa.

It is to be expected that many of the functional elements marked by accessible chromatin that are important for regulating gene expression in a given neural cell type would show dynamic accessibility across the lineage (i.e. stem cell specific enhancers would not be expected to be open in mature neurons). We examined regions of differential chromatin accessibility to determine the extent to which chromatin accessibility is changed during development of the nervous system. Hierarchical clustering of regions of chromatin with differential accessibility between cell types reveals two major clusters in which chromatin is either open in stem cells but inaccessible in neurons, or vice versa (Figure 3C). Intriguingly, there are other clusters where maximal chromatin accessibility is observed in either GMCs/ early neurons or larval neurons. Therefore, it is not as simple as NSC accessible regions progressively closing during differentiation and neuronal regions gradually opening. There are a large number of loci that are inaccessible in NSCs, then open in the intermediate GMCs/ newly born neurons stage before being rendered inaccessible again in terminally differentiated neurons (Figure 3C). In addition, a cluster enriched in larval neurons demonstrates that the chromatin accessibility landscape of larval neurons, although similar, is distinct from adult neurons.

Regions of open chromatin are thought to identify functional regulatory elements such as enhancers. Therefore, it is to be expected that these regions will be enriched for motifs belonging to transcription factors involved in neurogenesis. Identification of enriched motifs in sequences that were accessible in NSCs showed that expected transcription factor binding sites were highly enriched. For example, the E-box motif – CAGCNG – which is bound by the NSC proneural factor ase (Figure 3D) [21, 22]. Regions in which open chromatin was specifically enriched in mature neurons yielded novel sequence motifs for which we could not identify known transcription factors.

Gene ontology (GO) analysis of genes at which enriched chromatin accessibility was observed yielded expected biological process terms for each of the cell types examined. For example, terms such as "neuroblast fate determination and "chromosome segregation were enriched in stem cells relative to neurons, whilst "regulation of behaviour and "synaptic vesicle docking during exocytosis were enriched for differentiated neurons but not NSCs (Fig 3E).

### Chromatin accessibility in adult midgut cell types

Having observed chromatin accessibility changes in the cells of the developing CNS, we asked whether similar patterns would be observed in adult somatic stem cell lineages. The Drosophila midgut contains a pool of cycling intestinal stem cells (ISCs) that persists in the adult to maintain a population of terminally differentiated cells which mediate the absorptive and secretory functions of the organ [23, 24]. In contrast to neurogenesis, a single committed immature progenitor cell (enteroblast – EB) is produced from stem cell divisions, which then differentiates without further divisions to produce the mature epithelial cells of the midgut [25]. To examine chromatin accessibility in the cells of the adult midgut, we expressed *Dam* in the ISCs and EBs, as well as in the terminally differentiated absorptive cells, the enterocytes (ECs)(Fig 4A).

As with the CNS data, we observed predictable changes in chromatin accessibility at loci for genes with variable expression in the lineage. For example, *escargot* (*esg*) a transcription factor required for ISC self-renewal [26], displays multiple peaks of accessible chromatin at the gene body and surrounding region in ISCs and EBs, whilst little signal is observed in the ECs (Fig 4B). In contrast the *nubbin* locus (encoding EC marker – pdm1), displays peaks predominantly in the EC data, with relatively closed chromatin in the progenitor cell types (Fig 4C). As previously, hierarchical clustering revealed two major groups in which accessible chromatin was unregulated in either ISCs or ECs (Figure 4D). Smaller clusters were again evident in which accessible chromatin was up or downregulated exclusively in the intermediate EBs. However, this was much less pronounced than the changes observed in GMCs/early neurons of the developing CNS. This indicates that, similar to the CNS lineages, the majority of chromatin accessibility changes involved in specifying the fully differentiated cells do not occur until after EB maturation.

ISCs and NSCs fulfil similar roles in their respective organs in the production of highly specialised functional cells. However, whilst NSCs exist for a short amount of time during fly development to produce relatively long-lived neurons that persist in the adult CNS for the animal’s lifetime, the ISCs act post-developmentally to constantly replenish ECs in the adult gut. By comparing the chromatin accessibility of these two cell types, it is apparent that there are similarities in their chromatin states. Given the similarities that we observed for individual loci between CNS and midgut lineages, we queried whether it was possible to observe trends between the cells in the two lineages on a global scale. Principle component analysis reveals two distinct clusters for the two lineages in which >80% of the variance is explained in the first two principle components (Fig 4E). By examining the overall correlation between all cell types we observed a number of interesting features. Firstly, as expected all cell types correlated most closely with either their direct progeny or progenitor cell (Figure 4F). Therefore by clustering the data we were able to recapitulate the familial relationship between the cell types of the two lineages. The greatest similarities were observed between the intermediate progenitors and their cognate stem cells (R^2^ = 0.94 / 0.98 for CNS and midgut respectively). Interestingly, the greatest correlation outside of a lineage was between the two stem cell types (R2=0.76), whilst differentiated cells exhibited only weak correlation (ISCs vs NSC, R2 = 0.51). This indicates that somatic stem cell types may utilise a broadly similar chromatin landscape for the maintenance of multipotency, whilst lineage specific variation is relatively small.

**Figure 4.**
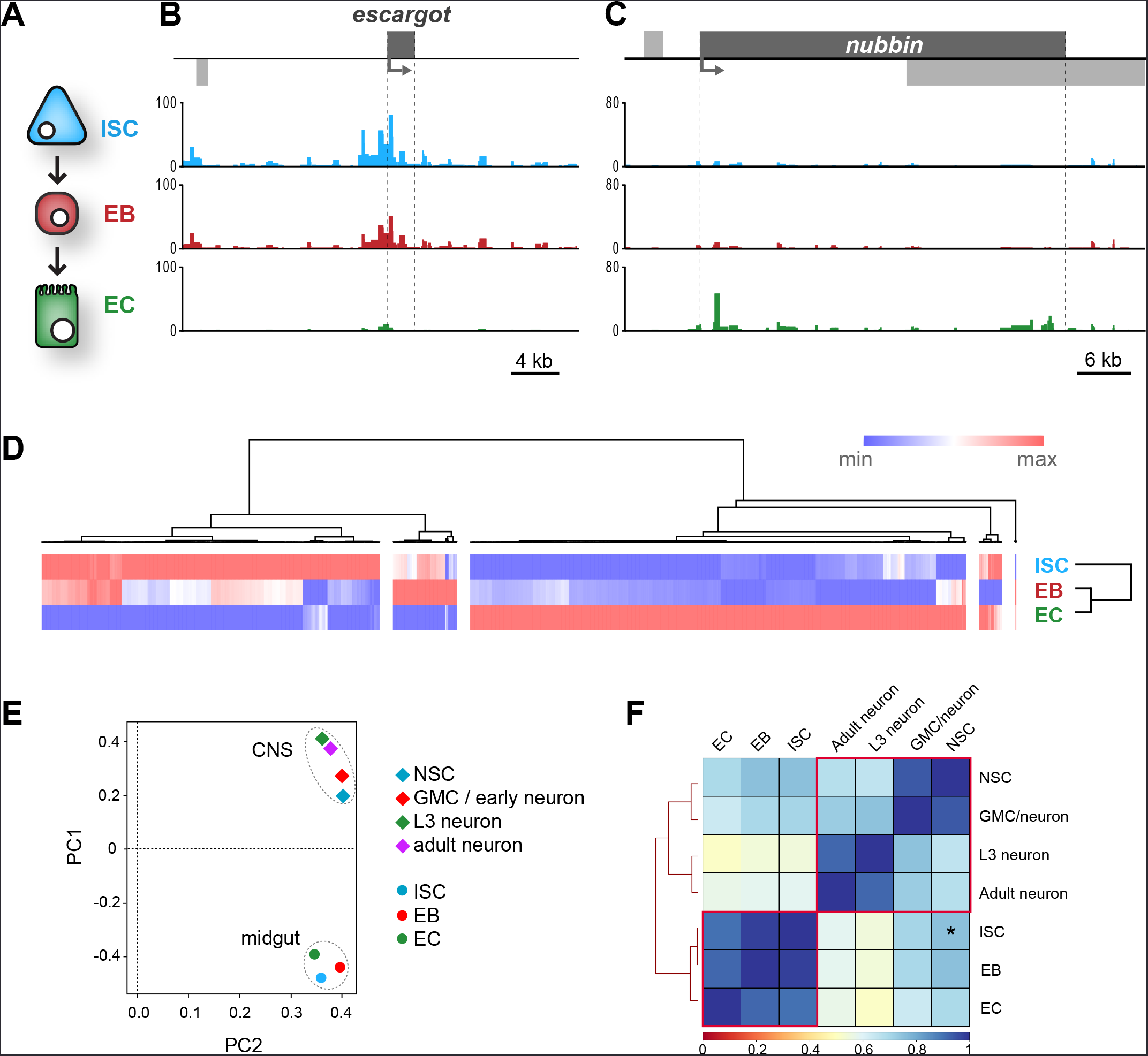
Dam chromatin accessibility profiling of cells in the adult midgut. **A)** Schematic of midgut lineage progression indicating cell types examined in this study. **B)** Chromatin accessibility displays expected trends at the *escargot* loci, known to be expressed exclusively in ISCs and EBs, but not ECs. Upstream promoter region shows greatest chromatin accessibility in ISCs, compared to other cell types. Similarly, dynamic peaks are observed in both 3’ and 5’ distal regions (putative enhancer regions), which are absent in ECs. y-axes = reads per million (rpm). **C)** Chromatin accessibility at the *nubbin* loci, known to be expressed exclusively in ECs. y-axes = reads per million (rpm). **D)** Hierarchical clustering of differentially accessible regions in gut cell types. Major clusters are observed in which accessible chromatin is enriched specifically in either ISCs or ECs, whilst smaller clusters indicate fewer regions with up or down-regulated accessibility in EBs. **E)** Principle component analysis (mean of all replicates) indicates distinct groupings of both lineages. **F)** Correlation matrix (Spearman’s rank) of means of all cells in CNS and midgut lineages. Individual lineages denoted with red outline. Note relatively high correlation between NSC and ISC (Asterisk – R2 = 0.76), whilst NSC correlation with EC and adult neurons are comparable.

### Enhancer prediction from Dam Accessibility data

Enhancer activity is closely linked to gene expression, therefore many tissue specific enhancers are required to orchestrate correct spatial and temporal transcription [27]. However, identification of functional enhancers can be challenging. Chromatin accessibility data has previously been used to identify novel enhancers [16, 28]. We reasoned that it would be possible to identify genomic regions corresponding to cell-type specific enhancers by comparing dynamically accessible regions between cell types. In support of this, we observed that the sequence covered by the *71C09-GAL4* line used in this study to profile GMCs/newly born neurons, displayed a higher peak specifically at this region than in either the stem cell or differentiated neuron data (Supplementary figure 4). Interestingly, a clear peak can still be observed in the NSC data, without concomitant reporter expression. Therefore, an enrichment of accessible chromatin does not necessarily correspond to an active enhancer in a given cell type. This is consistent with previous observations that DNase hypersensitive regions are often not active enhancers [29, 30].

We selected accessible regions with large differences between at least two cell types in the lineage, which satisfied various criteria for us to designate them as putative enhancers (see methods). We then identified available reporter lines from the Vienna tiles (VT)[31] and FlyLight [32] collections of GAL4 reporter lines that contained sequences encompassing our predicted enhancers upstream of a GAL4 reporter, and verified their expression in the tissues of interest. We identified enhancer-GAL4 lines in which reporter expression matched our predictions for enhancer activity. In the CNS Vienna line VT017417 and FlyLight line GMR56E07 both showed expression in the early part of the lineage in the CNS, with GFP reporter expression detectable predominantly in NSCs and GMCs (Fig5-A,B). This is consistent with accessible chromatin readings from our CATaDa data for these cell types in which progenitor cells displayed prominent peaks, whereas differentiated neurons did not. Similarly, we were able to detect functional cell-type specific enhancers in the midgut. The Vienna line, VT004241, showed reporter expression predominantly in *Delta* positive ISCs (Fig 5C). Therefore, it is possible to use CATaDa data to identify novel cell-type specific enhancers in multiple tissues.

**Figure 5.**
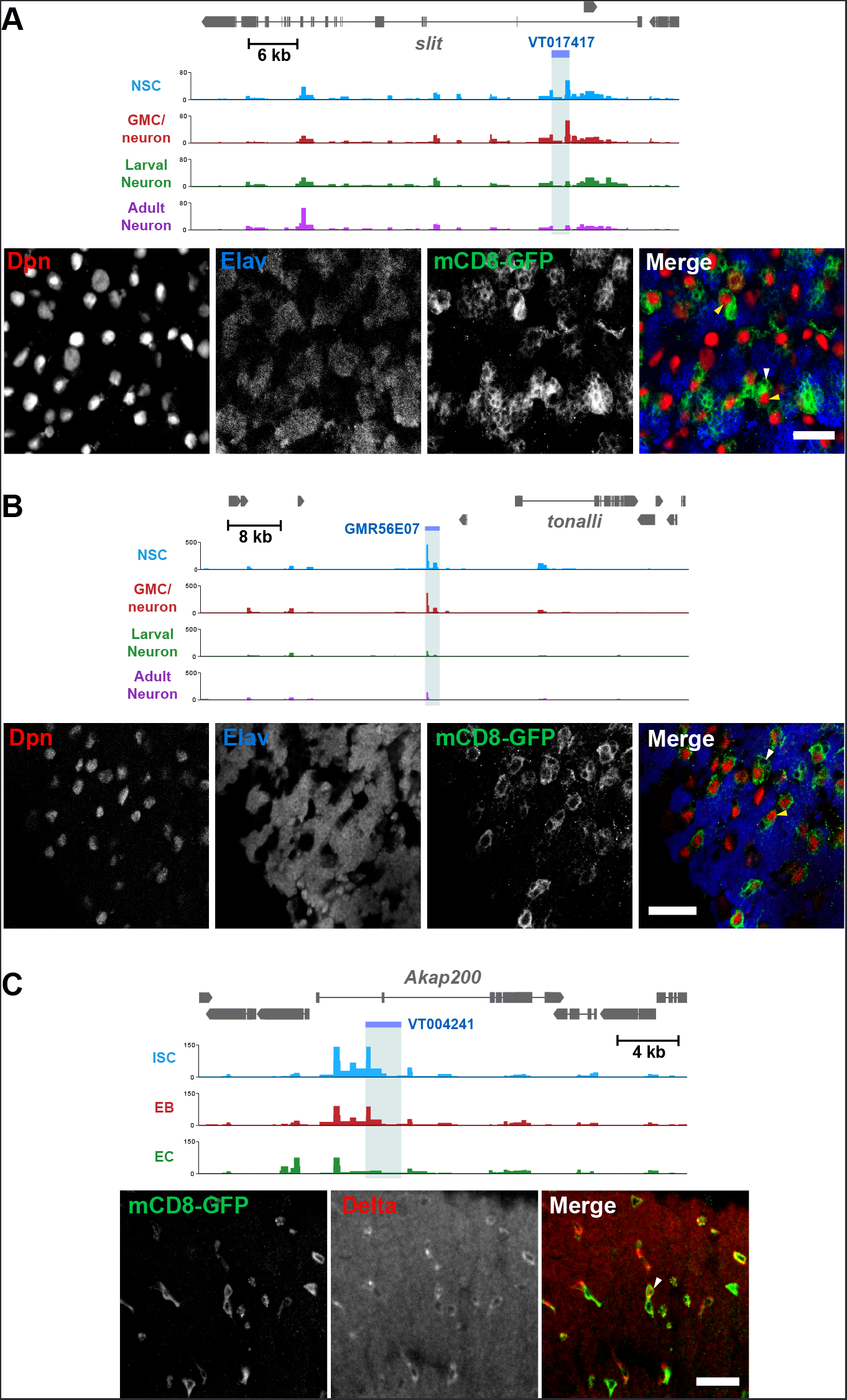
**Identification of cell-type specific enhancers from Dam accessibility data. A)** Intronic putative enhancer region VT017417 within *slit* locus reveals expression of GFP reporter gene predominantly in GMCs (white arrow) and newly born neurons, as well as some NSCs (Dpn positive, yellow arrow). **B)** Intergenic putative enhancer region GMR56E07 shows expression of GFP reporter gene predominantly in NSCs (yellow arrow), with some GMC expression (white arrow). **C)** Intronic putative enhancer region VT004241 Shows expression of GFP reporter predominantly in the ISCs (marked with Delta, white arrow). All scale bars = 20 µm.

### Distribution of sequencing reads reveals changes in global chromatin accessibility during lineage development

It is commonly accepted that the global accessibility of chromatin in a given cell type correlates broadly with potency. Whilst there are multiple lines of evidence showing that pluripotent cells have high levels of open chromatin, little data exists to support this idea in somatic multipotent stem cells and their progeny. We reasoned that by examining the distribution of normalised mapped sequencing reads (for each GATC fragment) we would be able to determine the nature of open chromatin in a given cell type.

Visual inspection of read distributions indicated that differentiated neurons had fewer GATC fragments with lower read counts (Supplementary figure 5A). To increase the significance of our analysis of read distributions, we decided to incorporate further neuronal and NSC datasets into our analysis which were available to us as control data from existing DamID studies ([13] and unpublished data). Including our previously described neuronal data, we examined the distribution of twelve adult and larval neuronal Dam accessibility datasets (derived from individual post-mitotic neuronal subtypes, cholinergic, GABAergic, and glutamatergic) and compared to data derived from NSCs.

From these distributions it is apparent that majority of GATC fragments in the genome have very few corresponding mapped reads in all samples, whilst fragments having over ~10 reads per million (rpm) were relatively infrequent. In other words, most of the genome is inaccessible or accessible at very low levels, whilst hyper-accessible regions are comparatively rare in all cell types(Fig 6A). The greatest difference between the distributions of neurons and NSCs was apparent at very low read numbers. We observed that there was a significantly greater proportion of GATC fragments with low read counts but without being completely inaccessible (~1-3rpm), in the NSC data compared to neurons (Fig 6A). This abundance of genomic regions in the stem cells with low-level chromatin accessibility indicates that open chromatin is more broadly distributed than in NSCs than neurons. We find that this trend is also apparent in the intermediate progenitor cell types, having intermediate amounts of GATC fragments mapping to low read counts (Figure 6B, supplementary figure 6A). Conversely, we observed a trend towards greater number of GATC fragments to which zero reads were mapped as differentiation progresses (Supplementary figure 6B). These data demonstrate that differentiated cells are more likely to have regions of chromatin that are completely inaccessible to Dam, indicating a globally lower amount of accessible chromatin in neurons. These trends are also observed in the midgut cells, implying that global changes to accessible chromatin are a common feature of somatic stem cell lineages *in vivo* (Supplementary figure 6C).

**Figure 6.**
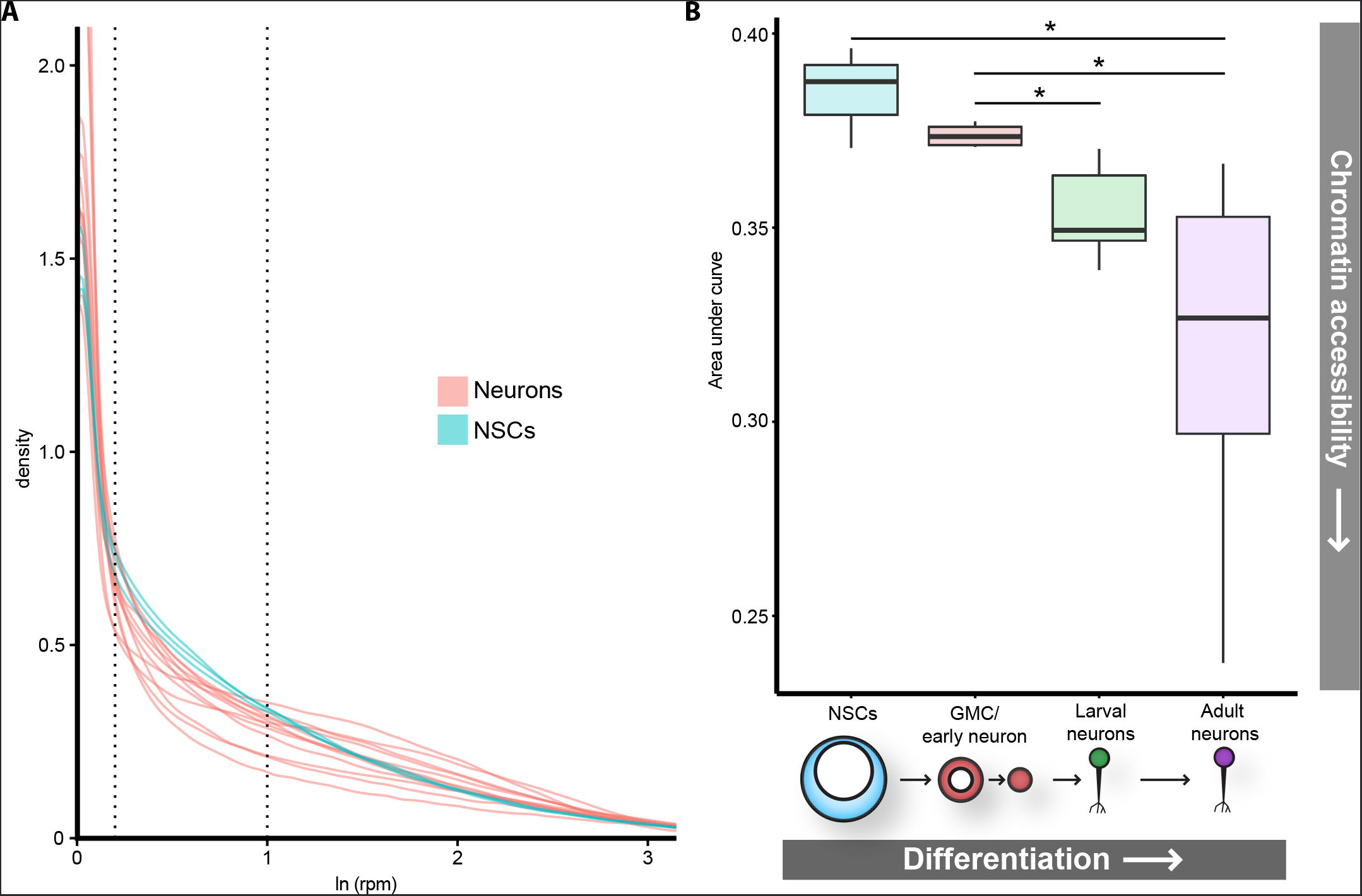
**Global chromatin accessibility is reduced in differentiated neurons. A)** Log transformed distribution of read counts at GATC fragments for neuronal (pink) or NSC replicates (blue). In addition to adult neuron data described in previous figures, CATaDa data for cholinergic, glutamatergic, and GABAergic adult neurons are included. NSC data includes extra replicate from [13]. **B)** Areas under curve for region bound by dotted lines in (**A**). Corresponding to ~1-3rpm. Data includes extra neuronal and NSC replicates shown in (**A**), as well as corresponding replicates at L3 for gutamatergic, GABAergic, and cholinergic neurons. Note that area under curve corresponds to proportion of GATC fragments having mapped reads within indicated range (i.e. NSCs have ~38% (median) of all GATC fragments within 1-3rpm mapped reads, compared to ~32% for adult neurons. Results were considered significant at \**P <* 0.05.

## Discussion

Recent studies have provided insights into chromatin accessibility of individual cell-types using accessibility assays coupled with cell sorting [33]. Whilst these strategies have been proven to produce meaningful biological data, they suffer from being technically challenging, particularly with regards to cell isolation. We have demonstrated that CATaDa yields chromatin accessibility profiles for defined cell types in vivo without the need for cell isolation, fixation or the extraction of naked chromatin. In addition to its ease of use, CATaDa also has the advantage that the marking of accessible DNA occurs in vivo. Due to this, there are no artefacts associated with cell disruption, chemical fixation or washing of the chromatin prior to the assay [34]. In addition, as Dam is expressed in vivo over several hours, the profiles produced will reflect dynamic changes to chromatin structure over the entire time period during which Dam is expressed.

CATaDa is limited by its resolution, which is restricted by the frequency of GATC sites in the genome (median spacing of ~200 bp in *Drosophila*). However, this can be increased by using a modified Dam in conjunction with immunoprecipitation (Dam-IP)[35]. Due to the dependence of Dam for methylation of GATC sequences, biases may be observed at loci which are depleted for GATC. It is worth noting that extensive sequence biases have also been reported for DNAse and ATAC-seq [36]. Although single cell protocols have recently been developed for chromatin accessibility techniques [37, 38], for routine experiments it is more usual to require a relatively large number of cells. For example for FAIRE-seq it is recommended to have a minimum of 1x10^6^ cells [1, 39], whilst DNase-seq typically requires 1x10^7^ cells [1, 40]. In contrast, DamID experiments can be performed with as few as 1000cells [41], therefore CATaDa is likely to also be effective with low cell numbers, making it competitive with ATAC-seq (500-50,000 cells) [7, 13]. Furthermore, single cell DamID has also recently been demonstrated, indicating that the minimum number of cells required for CATaDa is one [42].

Targeted DamID is rapidly being embraced by the *Drosophila* community with, at present, over 135 laboratories having requested the reagents and a number of papers already published ([15, 43-45]). Progress is also being made in adapting it for use in vertebrate models [41]. Whenever the binding of a protein of interest (POI) is investigated with this technique, Dam-only data (representing chromatin accessibility) for the cell type being assayed is also generated, as it is the control for which the Dam-POI is normalised. Therefore, researchers performing DamID experiments can now take advantage of this data, getting a “2-for-1 deal” whenever they use TaDa to profile the binding of a POI. Furthermore, much of this data is already available from published studies that could be readily analysed to provide novel biological insights.

The sequence of events which lead to repression of open chromatin in the transition between stem cells and their progeny is not well defined. Through identification of functional elements of the genome at various stages in this process, we can begin to understand how the dynamic chromatin landscape impacts the regulation and maintenance of differentiated cell states. Interestingly, we observe that chromatin accessibility in intermediate cell types is broadly more similar to their stem cell precursors than their differentiated progeny in CNS and midgut lineages (Figure 3B,C Figure 4B,C,F). This indicates that many stem cell specific regulatory regions remain accessible in intermediate cell types and are not fully repressed until terminal differentiation. This seems particularly surprising in the case of the EBs in the midgut considering that these cells do not undergo further mitotic divisions and are committed to a particular cell fate before their genesis by Notch signalling in the ISC [25]. Furthermore, in the intermediate cells of the larval CNS (GMCs and immature neurons), a relatively high proportion of cells profiled are neurons, which express markers thought to be associated with fully differentiated cells, suggesting that these regulatory regions may be open prior to terminal fate specification.

The reason for these regions of chromatin remaining accessible in stem cell progeny is unclear, however there are several plausible explanations. Firstly, regions that are bound by transcription factors that activate transcription may be replaced by transcriptional repressors. Such repressive factors are known to have detectable open chromatin “footprints”, similarly to activating factors [46]. Alternatively, the same factors that bind in the stem cells may recruit new binding partners that alter their activity to initiate repression rather than activation of gene expression. This explanation would require that further modifications occur to the chromatin following repressor activity as open chromatin regions are lost in fully differentiated cells, suggesting that repressors may no longer be bound. Retention of open chromatin regions in intermediate cell types may also reflect increased plasticity, indicating that cell fate has not yet been fully determined and that lineage reversion or dedifferentiation is possible given the introduction of the correct combination of factors. This idea is supported by the fact that immature post-mitotic neurons have been experimentally induced to dedifferentiate by interventions that are ineffective in fully differentiated adult cells [47]. It has been suggested that in some physiological contexts differentiated cells may revert to replenish stem cell pools [48, 49]. This idea may help to explain retention of plasticity in these cells types.

It is widely asserted that terminally differentiated cells have limited accessible chromatin whilst their progenitors maintain a broadly open chromatin landscape. However, there are few studies that investigate this phenomenon *in vivo*. Furthermore, the chromatin state in lineage committed intermediate progenitor cell types has been little studied. With CATaDa we have acquired evidence to indicate that NSCs in the developing *Drosophila* CNS appear to have more broadly open chromatin landscapes than their fully differentiated progeny. These differences are predominantly in the range of 1 to 3 rpm, which reflects chromatin with very low accessibility. Furthermore, the intermediate or immature progenitors, in both the CNS and midgut, retain a relatively open chromatin state, similar to that of their stem cells precursors (Figure 3B-C, Figure 4B,D,F). Together, these data support the model of stem cells containing more chromatin that has the potential to be accessed, whereas in differentiated cells, this flexibility is reduced and more regions are rendered completely inaccessible.

In conclusion, we have shown that cell-type specific chromatin accessibility profiles can be obtained through tightly controlled expression of Dam methylase. These data can be used to predict cell-type specific enhancers, as well as gaining insights into the global regulation of chromatin. Of particular interest are the dynamic changes in accessibility as cells progress towards terminal differentiation (e.g. the unique open regions observed in GMCs/early neurons) and the delayed compaction of stem cell gene associated chromatin. Also, analysis of the genome-wide distribution of chromatin accessibility supports a model of gradual compaction of large regions of low accessibility chromatin during differentiation. Overall, our results from profiling developing cell types illuminate the dynamic nature of chromatin accessibility in differentiation, and hint at organising principles which may apply to all somatic stem-cell lineages.

## Methods

### Fly stocks

*tub-GAL80^ts^*; *UAS-LT3-NDam* [15] was used to allow cell specific expression of *Dam*. The following *GAL4* driver lines were used to drive *Dam* expression in the CNS: *wor-GAL4* [50] for neuroblasts, *GMR71C09-GAL4* (Bloomington #39575) for GMCs and newly born neurons and *nSyb-GAL4* (Bloomington #51941) for mature larval and adult neurons. For expression of Dam in the gut the following lines were used for ISC, EB, and EC expression respectively: *esg-GAL4,UAS-2xEYFP/Cyo*; *Su(H)GBE-GAL80/TM3 Sb*, *Su(H) GBE-GAL4, UAS-CD8GFP/Cyo* and *{GawB}Myo31DF*^*NP0001*^*/CyO* [51, 52]. *P{tubP-GAL4}LL7/TM6*, *Tb* (Bloomington #5138) was used to drive ubiquitous expression in antennal-eye discs. *Cha^MI04508^-T2A-GAL4*, *Gad1^MI09277^-T2A-GAL4* and *vGlut^MI04979^-T2A-GAL4* driver lines were used to drive expression in cholinergic, GABAergic and glutamatergic neurons, respectively [53].

### Targeted DamID (TaDa)

To induce tissue specific *Dam* expression, *GAL4* driver lines were crossed to *GAL80*^ts^; *UAS-LT3-NDam* virgin females. Embryos were collected for 4 hours then raised at 18°C. Animals were transferred to 29°C at either 7 days after embryo deposition for 24 hours to obtain third instar larval tissues, or three days after eclosion to obtain adult heads or midguts. Fifty brains or thirty antennal-eye discs and midguts were dissected in PBS with 100 mM EDTA for each replicate. *71C09-GAL4* > *UAS-LT3-NDam* ventral nerve cords (VNCs) were dissected and central brain and optic lobe regions discarded due to presence of observed *71C09-GAL4* expression in a small subset of central brain neuroblasts.

Genomic DNA extraction and sequencing library preparation was performed as described previously [13], with minor modifications - MyTaq (Bioline) was used for PCR amplification of adapter ligated DNA . Libraries were sequenced using Illumina HiSeq single-end 50 bp sequencing. Two replicates of at least 10 million reads were acquired for each cell type. Sequencing data were mapped back to release 6.03 of the *Drosophila* genome using a previously described pipeline [54].

### Immunohistochemistry and imaging

*71C09-GAL4 > UAS-mCD8-GFP* third instar larval CNS or adult midgut were dissected in PBS and fixed for 20 min with 4% formaldehyde in PBS, 0.5mM EGTA, 5mM MgCl_2_. Tissues were stained with rat anti-elav (Developmental Studies Hybridoma Bank), chicken anti-GFP (Thermo scientific), mouse anti-Delta (Developmental Studies Hybridoma Bank), and guinea pig anti-deadpan (kind gift from A. Brand). Samples were imaged using a Zeiss LSM510 confocal microscope.

### Peak calling

Peaks were called and mapped to genes using a custom Perl program (available on request). In brief, a false discovery rate (FDR) was calculated for peaks (formed of two or more consecutive GATC fragments) for the individual replicates. Then each potential peak in the data was assigned a FDR. Any peaks with less than a 1% FDR were classified as significant. Significant peaks present in both replicates were used to form a final peak file. Any gene (genome release 6.11) within 5 kb of a peak (with no other genes in between) was identified as a potentially regulated gene.

### Motif identification

Regions of differentially accessible chromatin were identified by calculating GATC sites for which a difference of >10 rpm was observed between every replicate between at least two cell types. Hierarchical clustering was performed using Morpheus [55]. Sequences for major clusters showing enrichment for a given cell type were then analysed using MEME-ChIP [56].

### Gene ontology analysis

Potentially regulated genes, with a peak height of at least 10 rpm were submitted to GOToolBox [57] for GO analysis. GO term enrichments (frequency in data set divided by expected frequency) were calculated for each cell type.

### Cell type specific enhancer prediction

GATC fragments were identified with at least a 10 rpm difference in all replicates between at least two cell types. Any peak >2 kb from a transcriptional start site that did not overlap with coding sequence was designated an enhancer. Enhancers satisfying these criteria, which were covered by an available Vienna VT [31] or Janelia FlyLight [32] GAL4 line were chosen for validation based on the magnitude of the change in accessibility.

### Statistical analysis and data presentation

For comparing Dam data with ATAC and FAIRE, Monte Carlo experiments were performed using a custom Perl script (available on request). Comparisons of areas under curve in figure 5 were performed using Welch’s ANOVA for heteroscedasticity with Games-Howell post-hoc test in R. For data with equal variances, ANOVA was used with Tukey post-hoc testing (e.g. supplementary figure 6). Results were considered significant at \**P <* 0.05, \*\**P <* 0.01. Principle component analyses and correlation matrix plots were produced using deepTools [58]. Average TSS signal profiles were made using SeqPlots R/ Bioconductor package [59]. All other figures were produced using the ggplot2 package in R.

## Acknowledgments

We would like to thank Seth Cheetham, Jelle van den Ameele, and members of the Southall group for feedback and advice on this project. We would also like to thank Owen Marshall for his helpful discussions during the preparation of the manuscript and analysis of the data. We would like to thank Julia Cordero, Matthias Landgraf, Holly Ironfield and Benjamin White for providing fly stocks. Other stocks obtained from the Bloomington Drosophila Stock Center (NIH P40OD018537) and the Vienna Drosophila Resource Center (VDRC, <u>www.vdrc.at</u>). We also thank Andrea Brand for the Deadpan antibody. This work was funded by Wellcome Trust Investigator grant 104567 to T.D.S.

---

**Supplementary figure 1.**
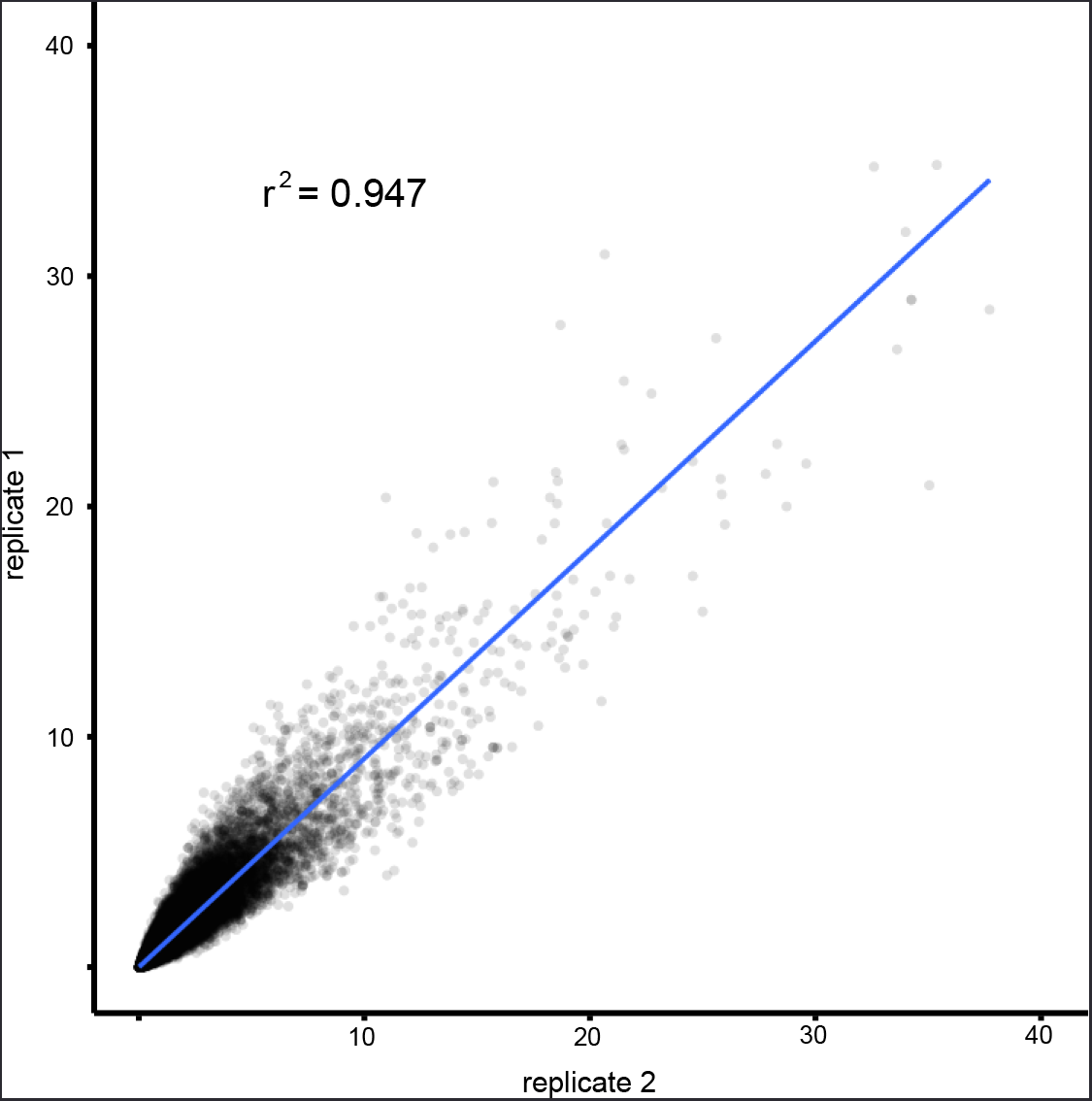
**Correlation between eye-disc Dam replicates**. Correlation between replicates shows good agreement indicating that the technique has high reproducibility. R^2^ = 0.94.

**Supplementary figure 2.**
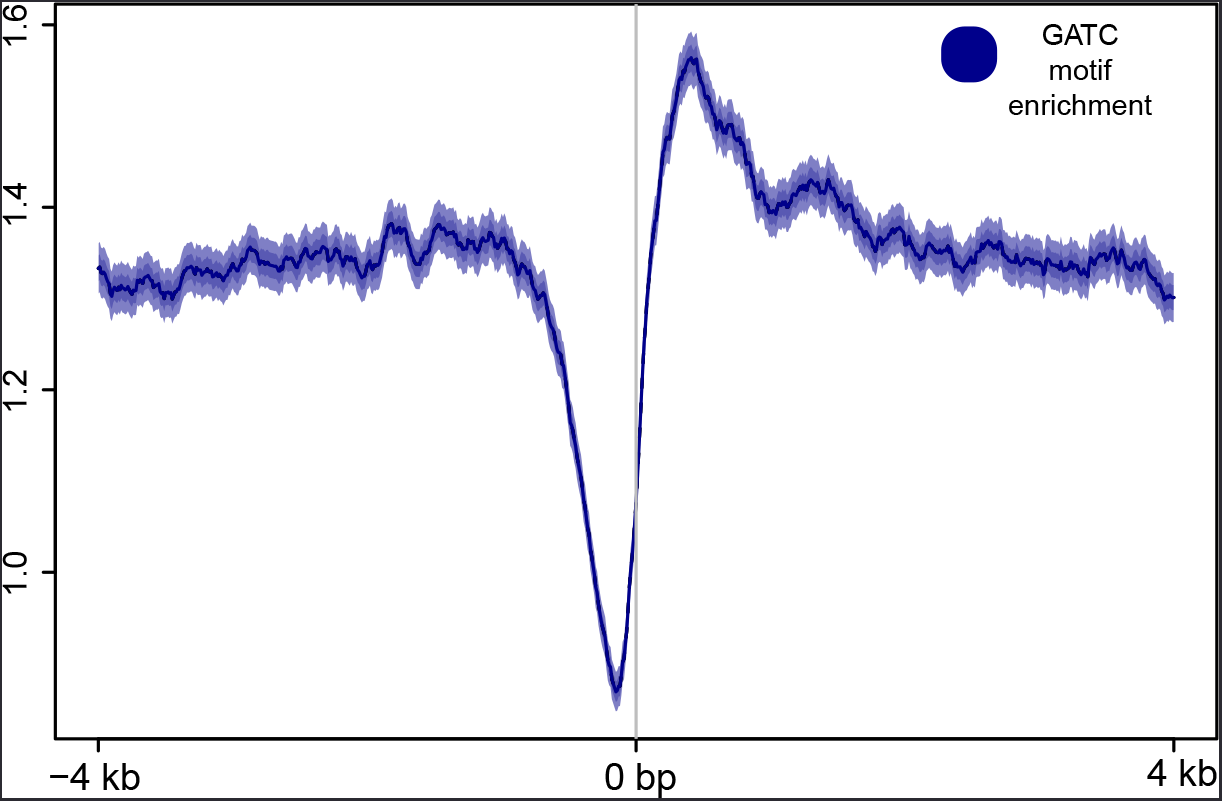
**Dam accessibility data shows reduced frequency of GATC sites at TSS**.

**Supplementary figure 3.**
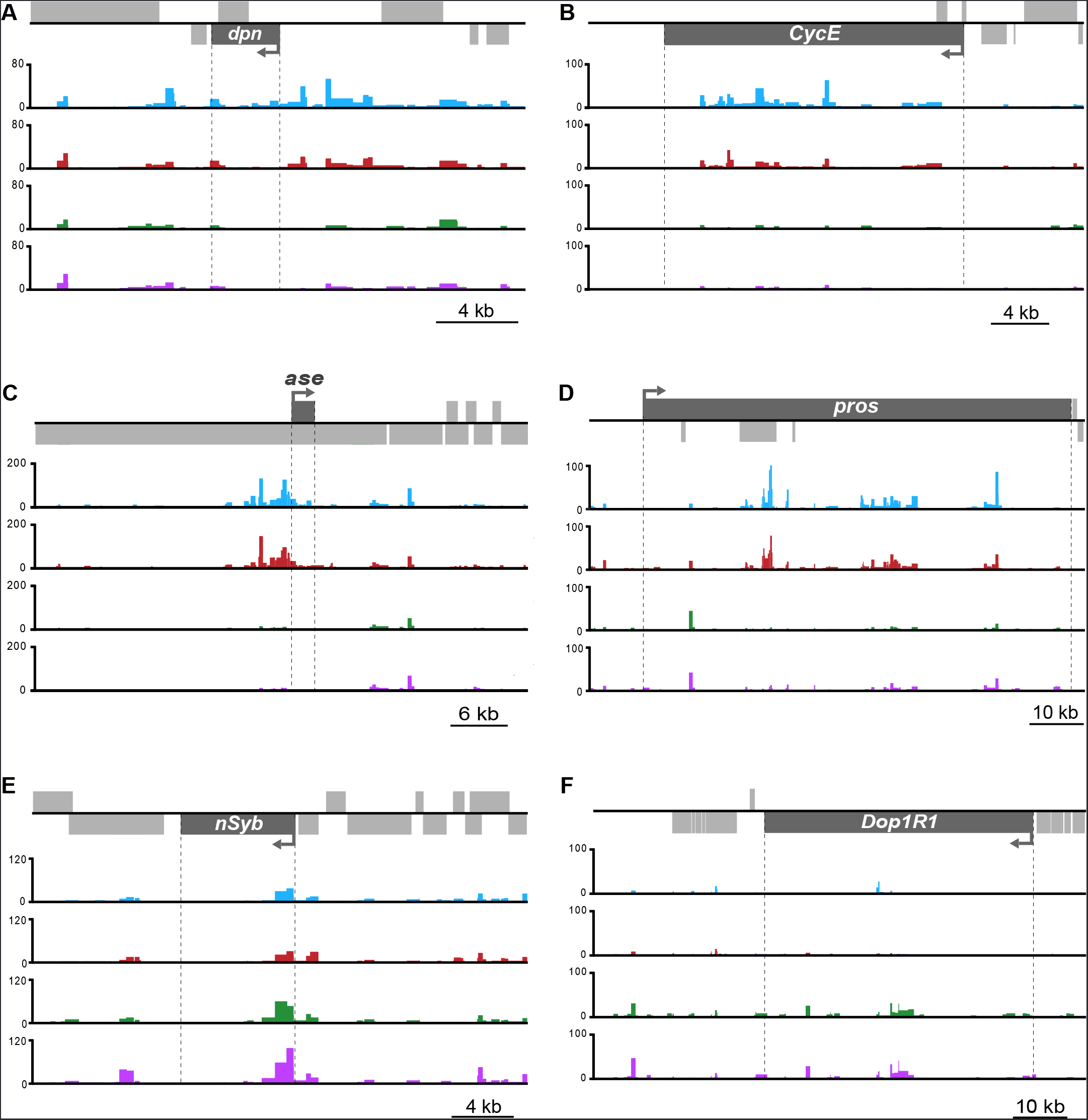
**Example loci showing dynamic chromatin accessibility in neuronal cell-types**. Regions of open chromatin are enriched in progenitor cell types for **A)** *deadpan* (*dpn*) **B)** *Cyclin E* (*CycE*) **C)** *asense* (*ase*), and **D)** *prospero (pros*); whilst peaks of greater accessibility are apparent in differentiated neurons at the **E)** *neuronal Synaptobrevin* (*nSyb*), and **F)** *Dopamine 1-like receptor 1* (*Dop1R1*) loci.

**Supplementary figure 4.**
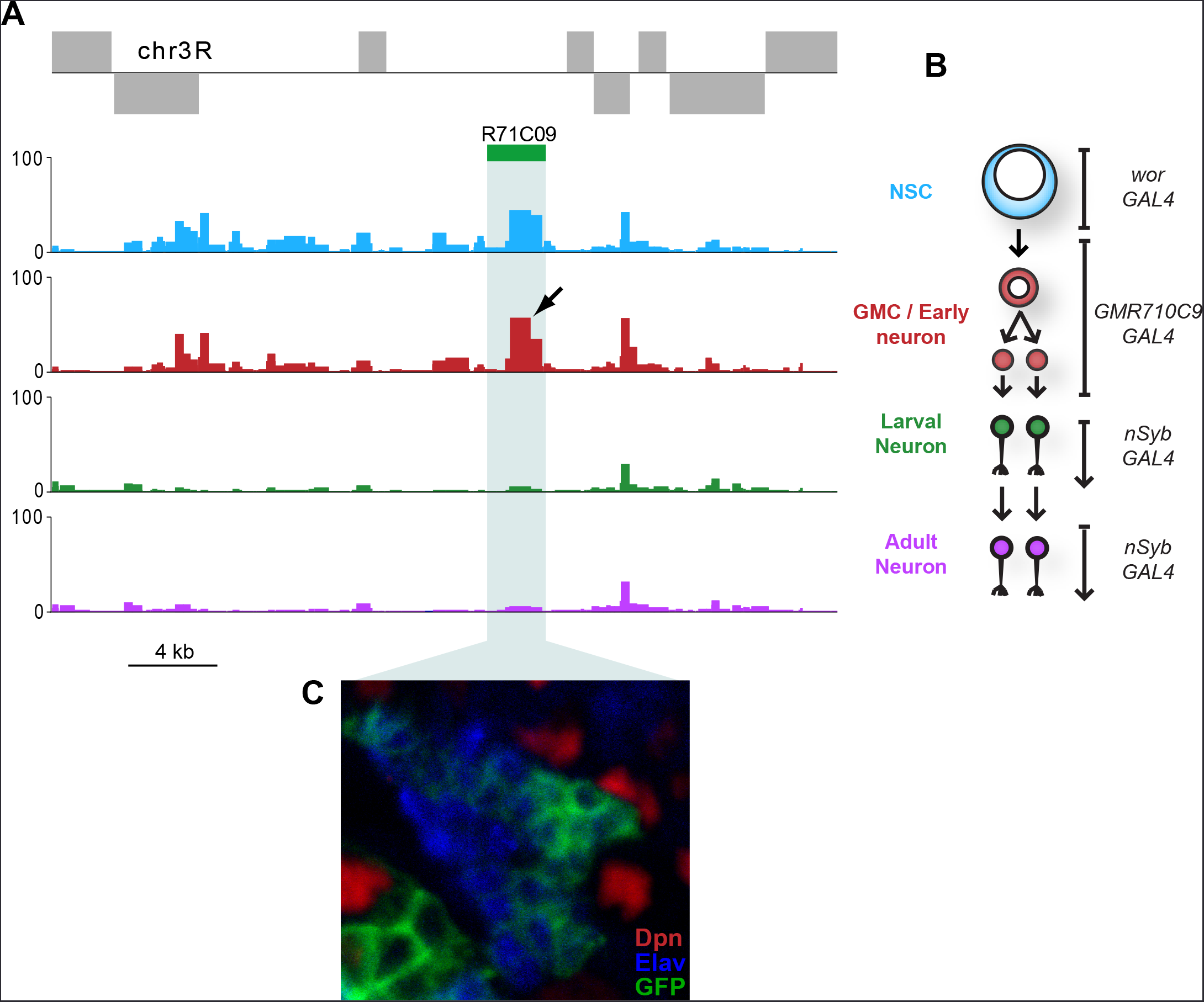
**Chromatin accessibility in neural cell types demonstrating dynamic accessibility of *R71C09* enhancer region - used to define GMC / immature neuron populations in this study. A)** A clear peak can be identified within the *R71C09* sequence which shows greatest accessibility in the GMCs and youngest populations of neurons in the lineage (arrow). **B)** Schematic indicating cell types assayed in this study with *GAL4* diver lines used to drive Dam expression. **C)** Expression pattern of *R71C09-GAL4/UAS-GFP* in the larval CNS. GFP is most strongly detected in cells adjacent to the neuroblast (Dpn - Red). These cells include presumptive GMCs (Dpn negative, Elav negative), and immature neurons (Elav - Blue)

**Supplementary figure 5.**
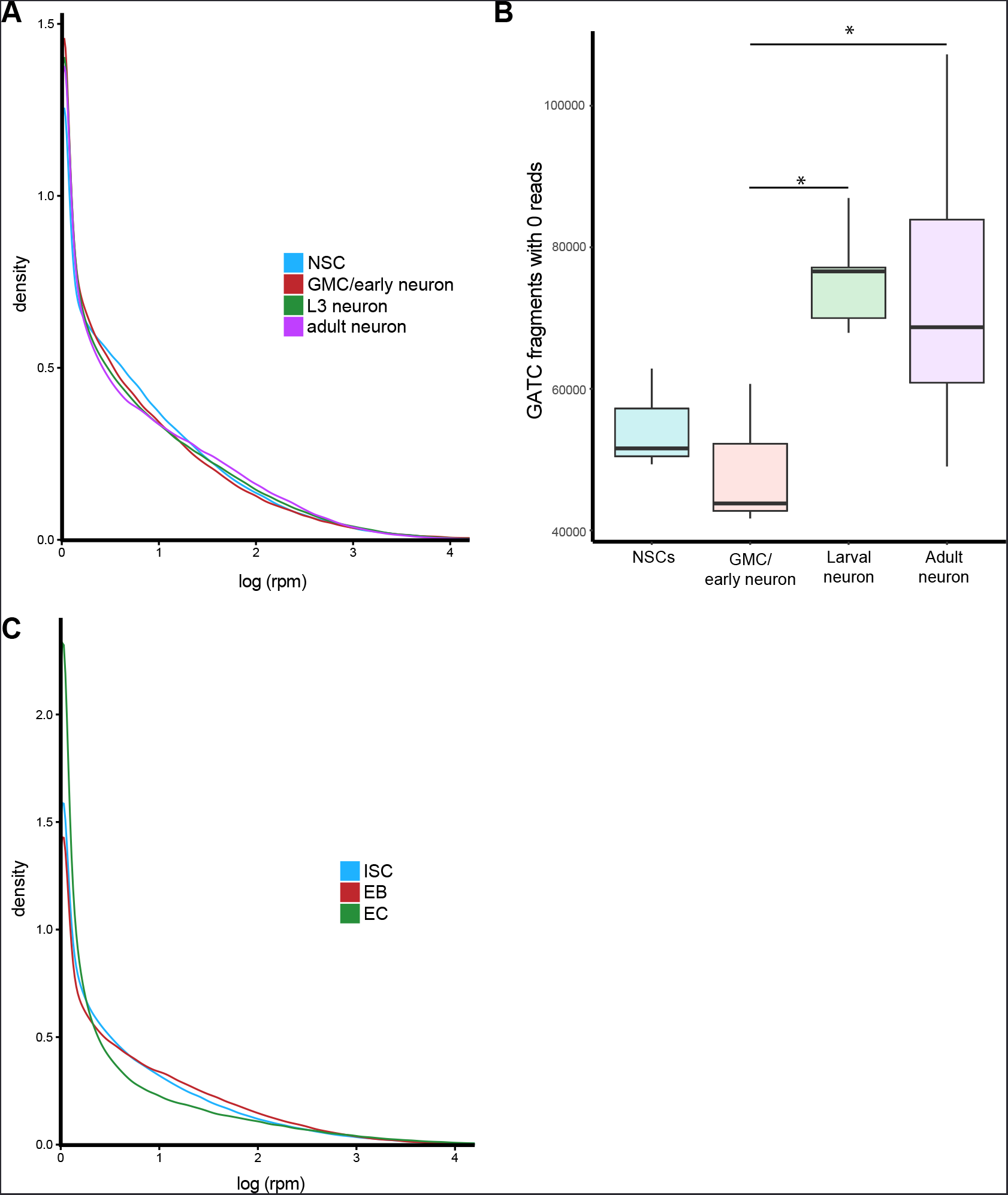
**Average distribution of sequencing reads for CNS and midgut cell types. A)** Mean distribution of reads for all cell types in the CNS. **B)** Number of regions with zero mapped reads for cell types of the CNS. **B)** Mean read distributions for all cell types in the CNS. Results were considered significant at \**P <* 0.05. **C)** Average distribution of reads for all cell types in the midgut.

